# Targeting ADORA-PDE10 cAMP Microdomain: A Novel Therapeutic Approach for Pulmonary Hypertension

**DOI:** 10.1101/2024.10.27.616460

**Authors:** Chanil Valasarajan, Golnaz Hesami, Giovanni Maroli, Anoop V. Cherian, Julio Castro-palomino Laria, Nahomi Castro-Palomino Laria, Astrid Wietelmann, Leoni Gnatzy-Feik, Stephan Rosenkranz, Rajkumar Savai, Werner Seeger, Soni Savai Pullamsetti

## Abstract

Despite substantial advancements in the treatment of pulmonary arterial hypertension (PAH), obstacles remain in achieving optimal outcomes. In this study, we focus on understanding the therapeutic effect of targeting the ADORA1/PDE10A-regulated cyclic AMP (cAMP) microenvironment in treating pulmonary hypertension. Screening of differentially expressed adenosine receptors in samples derived from healthy individuals and patients with idiopathic pulmonary arterial hypertension (IPAH) showed that ADORA1 expression was significantly upregulated under disease conditions. Functional studies revealed that, upon ADORA1 inhibition, donor hPASMCs showed anti-proliferative, pro-apoptotic features and increased intracellular cAMP levels. Surprisingly, the same effects were not replicated in IPAH PASMCs, suggesting the presence of another cAMP regulatory factor in close proximity to ADORA1 in IPAH PASMCs. To investigate this point, we performed protein-protein interaction studies in IPAH PASMCs and found that Phosphodiestrase 10A (PDE10A) co-localizes with ADORA1 under disease conditions. Moreover, we observed that ADORA1 and PDE10A form a regulatory complex in the A kinase anchoring protein 5 (AKAP5) microdomain. Silencing or dual inhibition of both ADORA1 and PDE10A in IPAH PASMCs induced anti-proliferative and pro-apoptotic effects with increased intracellular cAMP levels. To investigate the effects of the dual inhibitor in vivo, we administered it to both MCT and Sugen5416/hypoxia (SuHx) rat models of PAH, showing improved right ventricle function, reduced pulmonary vascular resistance and decreased lung vascular remodeling. In conclusion, we provide evidence that dual pharmacological targeting of ADORA1 and PDE10A has high therapeutic potential against PAH.

## Introduction

Pulmonary hypertension (PH) is a progressive, multifactorial disease of the pulmonary vasculature that is classified on the basis of clinical presentation, pathophysiologic mechanism, hemodynamic characteristics, and therapeutic management [1, 2]. Regardless of the clinical classifications, each group of PH exhibits pulmonary vasoconstriction, pathologic vascular remodeling, and right ventricular dilatation. The phenotypic changes seen in PAH (pulmonary arterial hypertension, or Group 1 PH) are mediated by various dysregulated pathway modulators such as transcription factors, growth factors, epigenetic modulators, metabolic regulators and non-coding RNAs [3]. Understanding these altered signaling pathways is crucial for exploring therapeutic options for the treatment of PH.

In PAH, the cyclic AMP (cAMP) signaling pathway has a significant impact on the pathogenesis of the disease, as the targeted use of various cAMP modulators to increase intracellular cAMP levels was shown to have beneficial effects [4, 5]. cAMP is a secondary messenger that helps mediating cellular responses to various extracellular stimuli. Intracellularly, cAMP levels are tightly regulated by two main enzymes –adenylyl cyclase (AC, enzymes catalyzing the synthesis of cAMP from ATP) and phosphodiesterases (PDEs, enzymes that degrade cAMP to 5′-AMP).

Among these cAMP-modulating enzymes, the PDE isoforms that degrade cAMP alone or both cAMP and cGMP are well studied with respect to the pathogenesis of PAH. Several PDEs exhibit differential expression and activity in both experimental and human PAH. PDE1, PDE3, PDE4, PDE5 and PDE10 are differentially regulated in the pulmonary vasculature of PAH [6], and PDE5 is targeted by sildenafil in an approved PAH therapy (Ref). However, PDE isoforms may be differentially distributed to induce localized cAMP regulation and specific signaling pathways, a phenomenon that remains unexplored in PH.

Recent studies have shown that cAMP signaling may be restricted to specific microdomains near the site of synthesis, leading to activation of nearby effectors [7]. The first indirect evidence of cAMP/PKA compartmentalization was observed in cardiac lysates by Corbin et al. in 1977 [8]. The non-uniform localization of cAMP signaling mediating proteins such as G protein-coupled receptors (GPCRs), ACs, PDEs, PKA and exchange proteins directly activated by cAMP (EPACs) suggests microdomain-specific signaling within cells [9, 10]. Studies have shown that the scaffold proteins such as A-kinase anchoring proteins (AKAPs) play an important role in mediating compartmentalized cAMP signaling [11]. Adenosine signaling is mediated by the activation of four GPCRs, namely adenosine receptor A1 (ADORA1), adenosine receptor A2a (ADORA2a), adenosine receptor A2b (ADORA2b) and adenosine receptor A3 (ADORA3) [12]. The adenosine receptors are mainly characterized on the basis of their adenosine requirement. For example, activation of ADORA1, ADORA2a and ADORA3 requires an adenosine concentration of 10nM-1μM, but ADORA2b requires a much higher adenosine concentration (10μM), which is rarely observed under physiological conditions [13].

Adenosine is a well-known pulmonary vasodilator that is often recommended as an alternative in pulmonary vasoreactivity testing to identify patients suitable for treatment with calcium channel blockers [14]. Importantly, circulating adenosine levels are low in patients with PAH, mainly due to increased adenosine deaminase activity [15]. However, one of the limiting factors for the use of adenosine as a therapeutic option in PAH is its extremely short half-life and adverse non-selective activation of multiple adenosine receptors [16]. Differential expression of adenosine receptors has been reported in different cell types of the lung under physiological conditions, mediating cell-specific signaling. For example, ADORA2b is abundantly expressed in bronchial SMCs, whereas ADORA1 and ADORA3 levels are low in these cells [17]. The spontaneous development of PAH in ADORA2a knockout mice suggests an important role of ADORA2a in the pathogenesis of PAH [18]. Moreover, the phenolic coumpound salidroside had beneficial effects in chronic hypoxia-induced PAH and reversed pulmonary artery remodeling by increasing the expression of ADORA2a and enhancing ADORA2a-mediated mitochondrial-dependent apoptosis [19]. On the other hand, activation of ADORA2b promotes the release of endothelin-1 and interleukin-6 from PAECs and PASMCs, and targeted deletion of ADORA2b has beneficial effects against PH development [20, 21].

Although several pharmacological treatments have been approved for PAH therapy, class-specific side effects have been reported and mortality remains high [22]. Thus, there is urgency to identify new therapeutic targets to address the unmet medical need in PH. The present study addresses the formation of a unique ADORA1-PDE10A complex stringently regulating the local cAMP levels in the AKAP5 microdomains under PAH conditions. The study also investigates the therapeutic potential of targeting the ADORA1-PDE10A complex using a dual inhibitor.

## Results

### ADORA1 expression is upregulated in IPAH PASMCs and correlates with reduced intracellular cAMP levels

To investigate the expression pattern of adenosine receptors in IPAH patients, we performed expression analysis at the mRNA and protein level in PASMCs and lung tissue, revealing upregulation of ADORA1 in IPAH versus donor samples (Fig. 1A, B). Immunofluorescence staining of ADORA1 in lung tissue also showed a similar expression pattern as previously observed in PASMCs (Fig. 1C). Previous studies have shown that ADORA1 negatively regulates the intracellular cAMP levels, thus we measured the intracellular cAMP levels in the IPAH and donor PASMCs, in line with the ADORA1 expression studies the levels of intracellular cAMP was decreased under disease conditions (Fig. 1D). These results suggest that ADORA1 can be a potential target to regulate intracellular cAMP level in the PAH patients.

**Figure 1.**
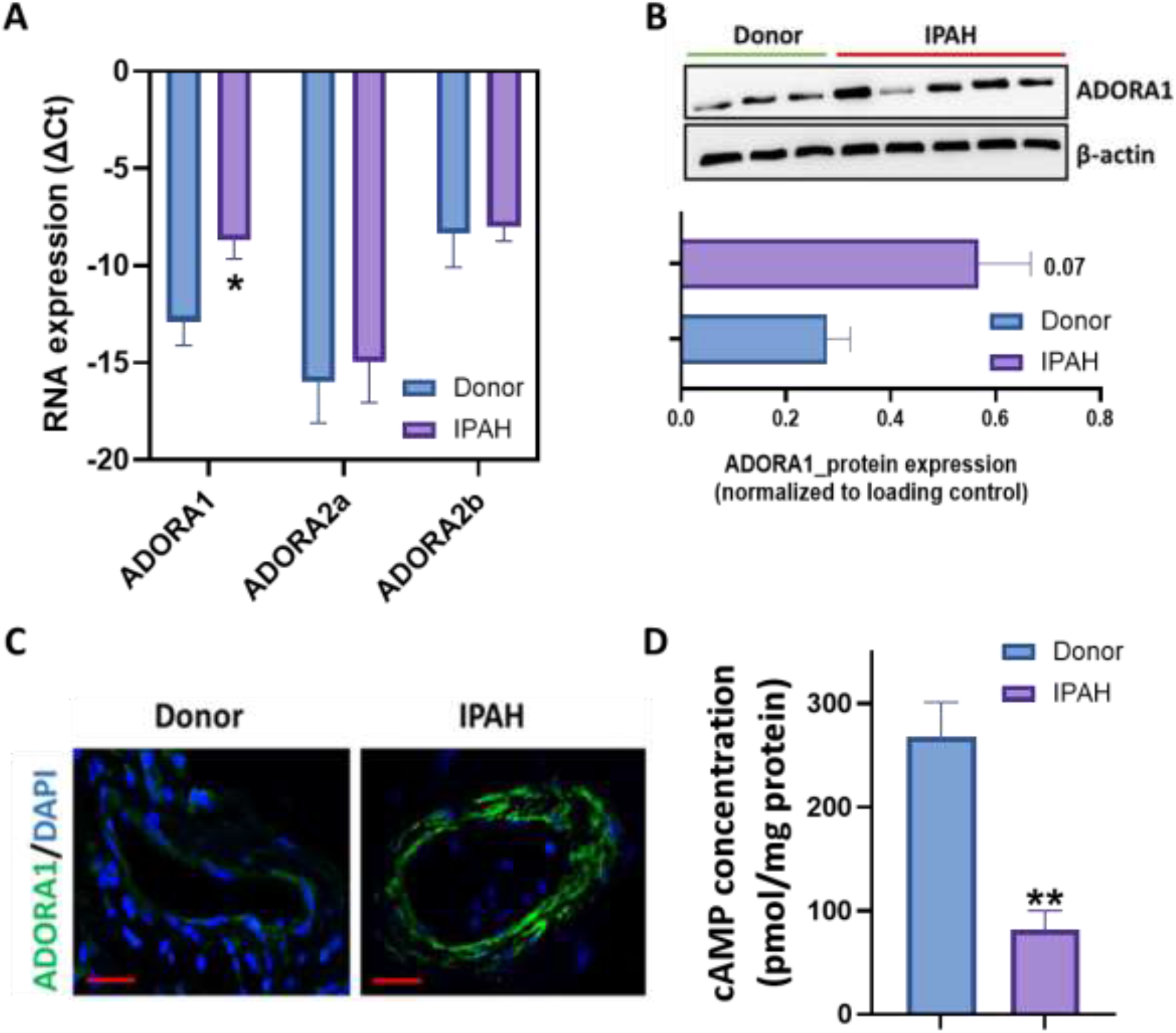
Expression analysis of adenosine receptors in donor and IPAH PASMCs. (A) Expression analysis of ADORA1, ADORA2a and ADORA2b by qRT-PCR between donor (n=3) and IPAH (n=3) PASMCs. (B) Expression analysis of ADORA1 by Western blotting between donor (n=3) and IPAH (n=5) PASMCs. (C) Immunofluorescence staining of ADORA1 in donor and IPAH lung tissues (n=3) scale bar=20μm. (D) Intracellular cAMP was measured using an EIA assay kit in donor and IPAH PASMCs (n=4). (H) Measurement of proliferation by BrdU assay in both donor and IPAH PASMCs, (n=3). (I) Measurement of apoptosis in donor and IPAH PASMCs using the cell death assay, (n=3). Bars show mean values ± S.E.M. *p < 0.05 & **p < 0.01, compared to donor PASMCs.

### ADORA1 silencing induces apoptosis in donor PASMCs but has no effect on the phenotype of IPAH PASMCs

To understand the functional significance of ADORA1 in PASMCs, we performed siRNA-mediated silencing of ADORA1 and antagonist-based inhibition of ADORA1 activity in healthy and IPAH PASMCs. Inhibition of ADORA1 induced apoptosis in donor-derived PASMCs without altering the proliferation rate (Fig. 2A,B,D,E). Intracellular cAMP levels were also increased upon ADORA1 inhibition in healthy PASMCs (Fig. 2C,F). Conversely, the effect observed in donor PASMCs was not replicated in PASMCs derived from IPAH patients (Fig.2 G-L). This observation prompted us to investigate whether ADORA1 forms a bidirectional regulatory complex to stringently maintain low intracellular cAMP levels upon PAH.

**Figure 2.**
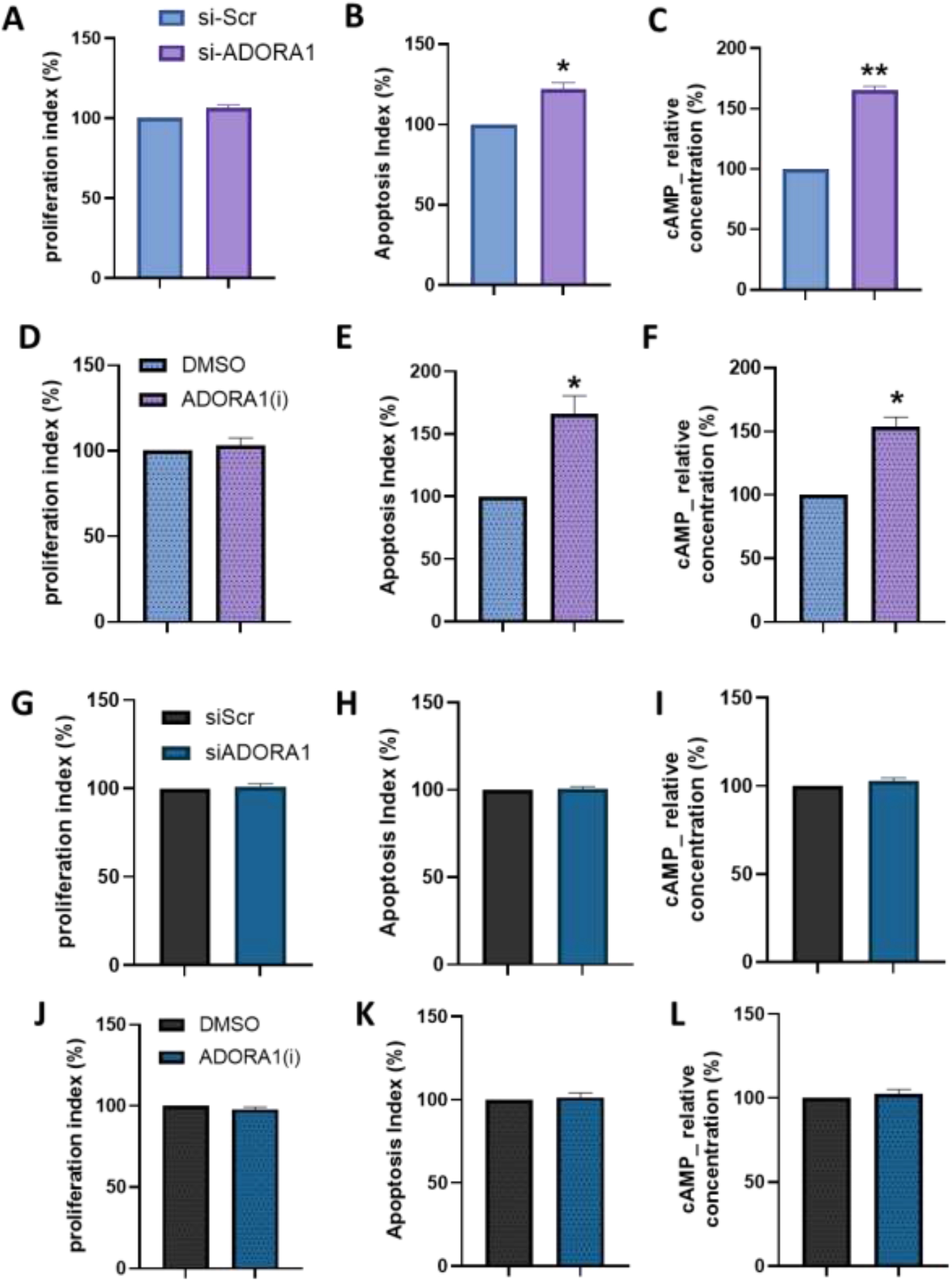
Functional effects of inhibiting ADORA1 in donor and IPAH hPASMCs. (A, D) Measurement of proliferation using BrdU assay (n=3), (B, E) measurement of apoptosis using the cell death assay kit (n=3), and (C, F) intracellular cAMP measurement using cAMP EIA kit (n=3) in donor PASMCS treated with siRNA and antagonist. (G, J) Measurement of proliferation using BrdU assay (n=3), (H, K) measurement of apoptosis using the cell death assay kit (n=3), and (I, L) intracellular cAMP measurement using cAMP EIA kit (n=3) in IPAH PASMCS treated with siRNA and antagonist. Bars indicate means ± S.E.M. Data were analyzed using paired t-test. *p <0.05, vs si-Scr /DMSO

### The ADORA1-PDE10A complex regulates cAMP levels in the AKAP5 microdomain under disease condition

Previous studies have shown that several cAMP-controlling PDEs were upregulated in patients with PH, leading us to hypothesize that ADORA1 might be in close proximity to a cAMP-hydrolyzing PDE under disease conditions (Fig. 3A). To search for the cAMP-hydrolyzing PDEs in close proximity to ADORA1, we performed immunocytochemical staining in IPAH PASMCs. Interestingly, we found that PDE10 co-localizes with ADORA1 (Fig. 3B,C), which was further verified by immunofluorescence in IPAH lung tissue (Fig. 3D). Previous studies have also shown that the expression of PDE10A is upregulated under disease conditions [23]. Furthermore, we performed co-immunoprecipitation assays for ADORA1 and PDE10A in healthy and IPAH PASMCs to validate the formation of a PAH-restricted complex. Strikingly, we found that ADORA1 and PDE10A form a complex only in PAH PASMCs, whereas this was not observed in healthy PASMCs (Fig. 3E). Moreover, we further confirmed the emergence of a PAH-specific complex with proximity ligation assays in healthy and IPAH PASMCs (Fig. 3F). Previous studies have shown that GPCRs and the PDEs regulate cAMP levels at the subcellular levels by forming microdomains with the help of anchoring proteins like AKAPs [24-26]. Interestingly, we found that the expression of AKAP5 was significantly upregulated in PAH versus donor PASMCs (Fig. 3G). Additionally, co-immunoprecipitation assays demonstrated that the ADORA1, PDE10A and AKAP5 form a supercomplex in IPAH PASMCs (Fig. 3H,I). These results show that ADORA1 and PDE10A form a bidirectional cAMP regulatory complex in the AKAP5 microdomain under disease conditions.

**Figure 3.**
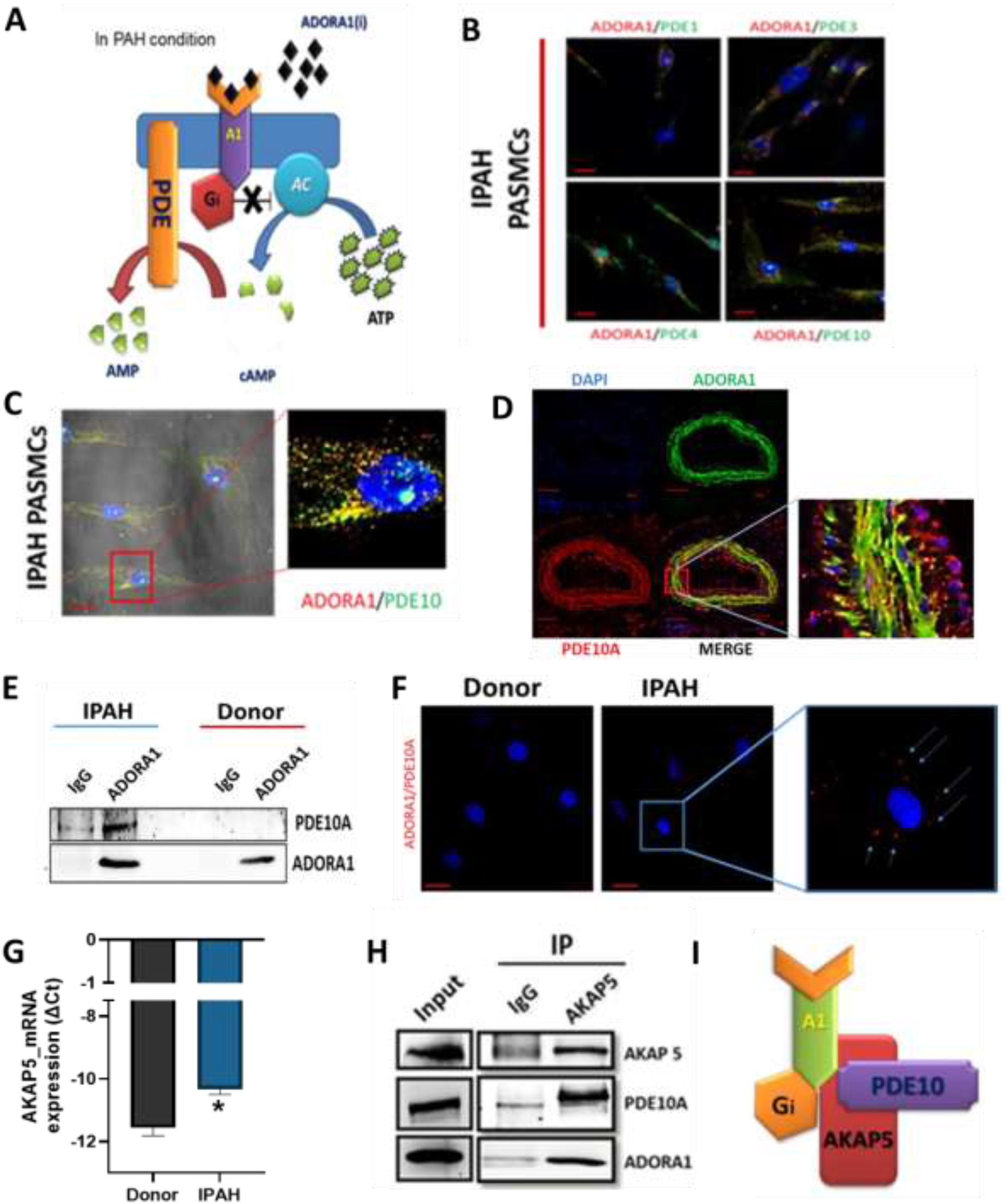
Formation of a cAMP bidirectional regulatiory complex under IPAH condition. (A) Schematic representation of hypothesized mechanism of cAMP regulation by ADORA1 and PDEs in IPAH PASMCs. (B) Immunostaining based screening of cAMP regulating PDEs (PDE1, PDE3, PDE4 and PDE10) co-localization with ADORA1 in IPAH PASMCs (n=3) scale bar= 20μm. (C) Confocal imaging of ADORA1 and PDE10A in IPAH PASMCs (n=3) scale bar= 20μm (D) Dual immunofluorescence staining of ADORA1 and PDE10A in lung tissue from IPAH patients (n=3) scale bar=50μm. (E) Co-IP was performed in IPAH- and donor-PASMCs by pulling down ADORA1 and blotting for both ADORA1 and PDE10A (n=3). (F) Confocal imaging of proximity ligation assay of ADORA1 and PDE10A in IPAH- and donor-PASMCs (n=3) scale bar= 20μm. (D) AKAP5 expression analysis using qRT-PCR between the donor-(n=3) and IPAH-PASMCs (n=3), (E) Co-IP in IPAH PASMCs by pulling down AKAP5 (n=3), (F) Representative image of the supercomplex. Bars indicate means ± S.E.M. Data were analyzed using the unpaired student t-test. **p <0.01 vs Donor.

### Dual inhibition of ADORA1 and PDE10A reverses the disease phenotype in IPAH-PASMCs

To test the effect of concomitant inhibition of ADORA1 and PDE10A, we performed simultaneous siRNA-mediated silencing of the two genes in IPAH PASMCs, which resulted in a strong anti-proliferative and pro-apoptotic phenotype in IPAH-PASMCs (Fig. 4A,B). Moreover, intracellular cAMP levels were strongly increased by concurrent knock-down of ADORA1 and PDE10A in IPAH PASMCs (Fig. 4C). Thus, we decided to investigate the therapeutic relevance of pharmacological inhibition of the ADORA1-PDE10A complex in PAH, by using a specific dual inhibitor (a single compound that inhibits both ADORA1 and PDE10A activity developed by Palbiofarma, Barcelona) in IPAH PASMCs. Similar to the siRNA-based strategy, treatment with the dual inhibitor reversed the pathologic phenotype of IPAH PASMCs (Fig. 4E,F) and significantly increased intracellular cAMP levels and downstream phosphorylation of cAMP response element-binding protein (CREB) in IPAH PASMCs (Fig. 4G,H). These results demonstrate that the genetic or pharmacological targeting of both ADORA1 and PDE10A reverses the pathological state of IPAH PASMCs in vitro, suggesting a possible therapeutic potential in PAH.

**Figure 4.**
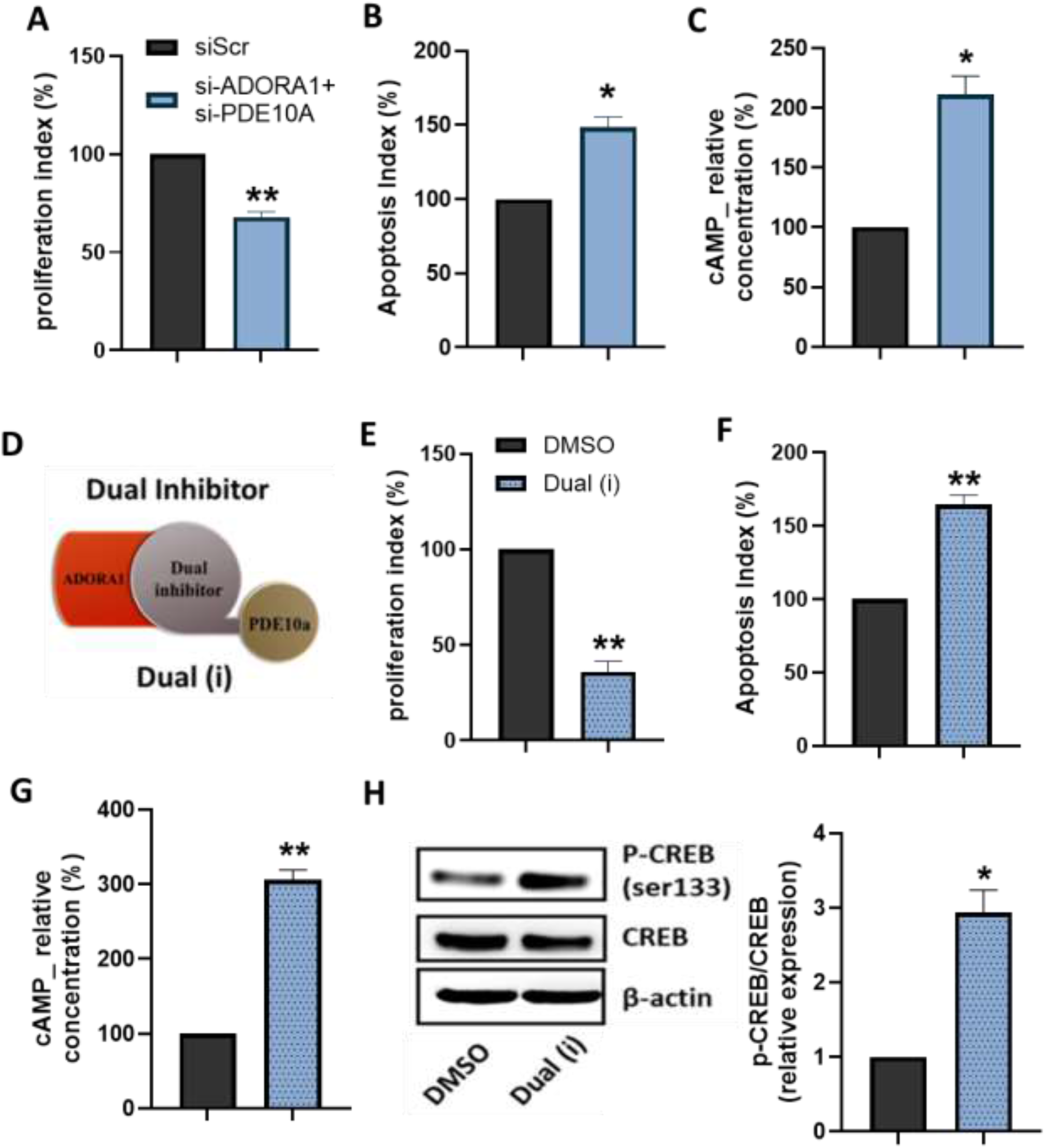
Effects of dual inhibition of ADORA1 and PDE10A in IPAH PASMCs. (A) Measurement of proliferation using BrdU assay (n=3), (B) measurement of apoptosis using the cell death assay kit (n=3), and (C) intracellular cAMP measurement using cAMP EIA kit (n=3) IPAH PASMCs after treatment using siRNA. (D) Representative image of dual inhibitor targeting ADORA1and PDE10A. (E) Measurement of proliferation using BrdU assay (n=3), (F) measurement of apoptosis using the cell death assay kit (n=3), and (G) Intracellular cAMP measurement using cAMP EIA kit (n=3) in IPAH PASMCs after treatment using dual inhibitor [dual(i)]. (H, I) Western blotting of p-CREB, CREB and (H, J) p27 in IPAH PASMCs treated with dual inhibitor. Bars indicate means ± S.E.M. Data were analyzed using paired t-test. *p <0.05, **p <0.01, ***p <0.001 vs siScr/ DMSO.

### Treatment with the ADORA1-PDE10A dual inhibitor improves cardio-pulmonary function in SuHx rats

We first investigated the therapeutic effect of the dual inhibitor in the Sugen5416-hypoxia rat model of PAH (Fig. 5A). Cardiac MRI measurements (Fig. 5B) showed that right ventricle (RV) ejection fraction (EF) was preserved in the dual inhibitor-treated group, while the placebo group showed progressive functional deterioration of the RV (Fig. 5C). Moreover, cardiac stroke volume (SV) and cardiac output (CO) improved significantly after treatment with the dual inhibitor compared to the placebo group (Fig. 5D,E), without significant changes in cardiac hypertrophy (Fig. 5F). Total pulmonary vascular resistance (TPVR) was calculated using RVSP measurement from right heart catheterization and cardiac MRI. We found that treatment with the dual inhibitor significantly reduced TPVR compared to placebo (Fig. 5G). Moreover, morphometric analysis of the lungs showed that the dual inhibitor-treated group had less vascular remodeling in terms of the number of muscularized pulmonary vessels and reduced medial wall thickness (Fig. 5H,I).

**Figure 5.**
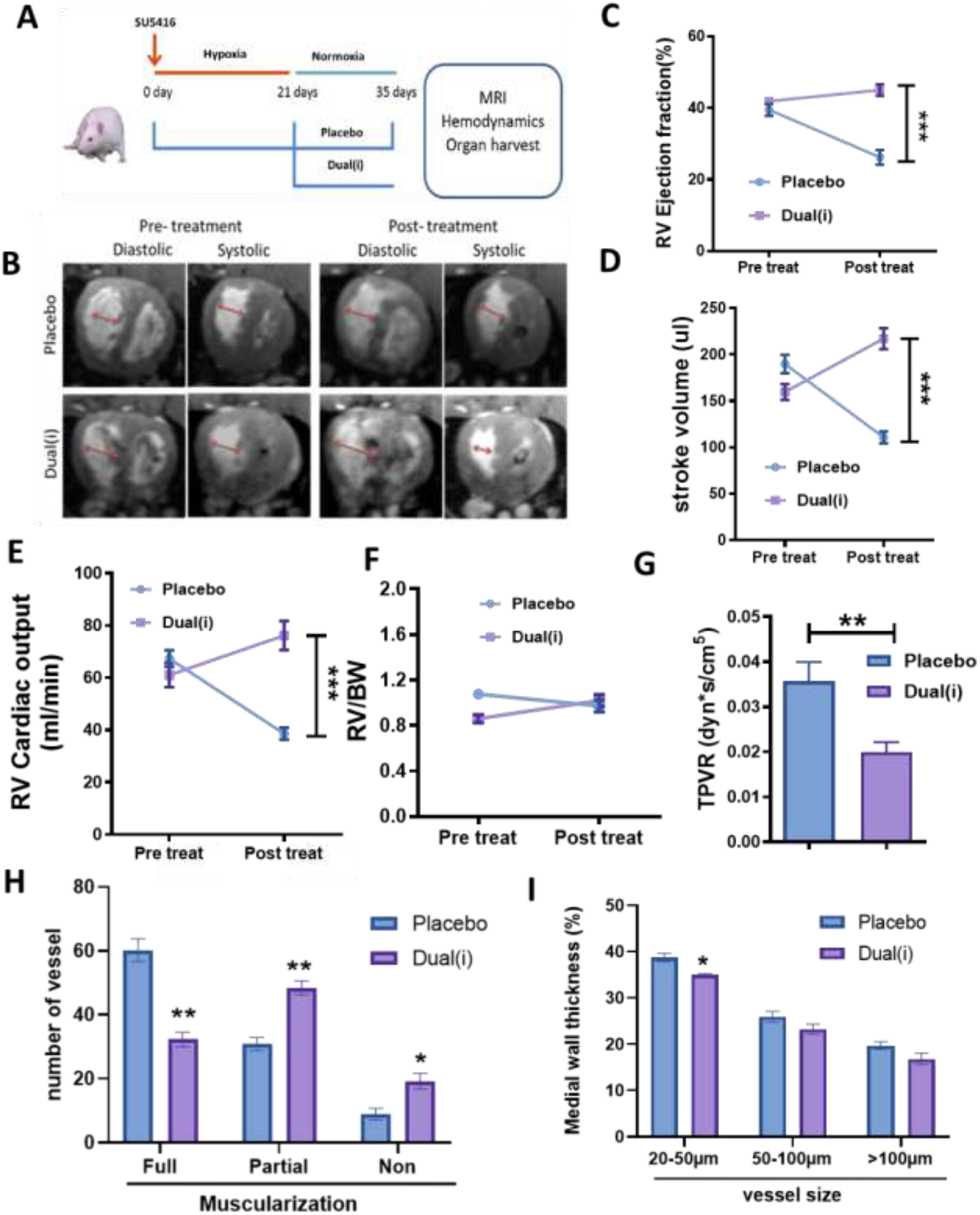
Effect of dual inhibitor on the SuHx-PAH rat model. (A) Schematic representation of SuHx-PAH rat model. (B) Cardiac MRI images of SuHx-PAH rats pre-and post-treatment with placebo or dual inhibitor. (C) RV ejection fraction, (D) RV stroke volume, (E) cardiac output, and (F) Fulton index (RV+LV+S) from cardiac MRI from Placebo (n=6) and dual inhibitor (n=6) treated groups. Bars indicate means ± S.E.M. Data were analyzed using two-way ANOVA with Tukey’s multiple comparison. ***p <0.001 vs Placebo. (G) TPVR calculated from RVSP and Cardiac output of placebo (n=6) and Dual inhibitor treated groups (n=6). Bars indicate means ± S.E.M. Data were analyzed using the unpaired student t-test. *p <0.05 *vs* Placebo. (H) Medial wall thickness measurements in pulmonary vessels, (I) muscularization measurements from placebo (n=4) and Dual inhibitor treated groups (n=4). Bars indicate means ± S.E.M. Data were analyzed using two-way ANOVA with Tukey’s multiple comparison. *p <0.05, **p <0.01, ***p <0.001 *vs* Placebo.

### Dual inhibitor treatment improves survival and RV function in the MCT rat model of PAH

The therapeutic efficacy of dual inhibition of ADORA1 and PDE10A was also investigated in a second rat PAH model induced by monocrotaline (MCT) treatment (Fig. 6A). Rats treated with the dual inhibitor showed better survival rate compared to placebo-treated rats (Fig. 6B). Moreover, treatment with the dual inhibitor improved cardiac function of MCT-treated rats in terms of RV ejection fraction, stroke volume and cardiac output (Fig. 6 D-F). Right ventricular hypertrophy assessed with the Fulton index was also significantly reduced in rats treated with the dual inhibitor compared to placebo (Fig. 6G). Finally, TPVR and vascular remodeling were also reduced with dual inhibitor treatment (Fig. 6 H-J). These results show that inhibition of ADORA1 and PDE10A by a dual pharmacological inhibitor has a therapeutic effect in two different rat models of PAH.

**Figure 6.**
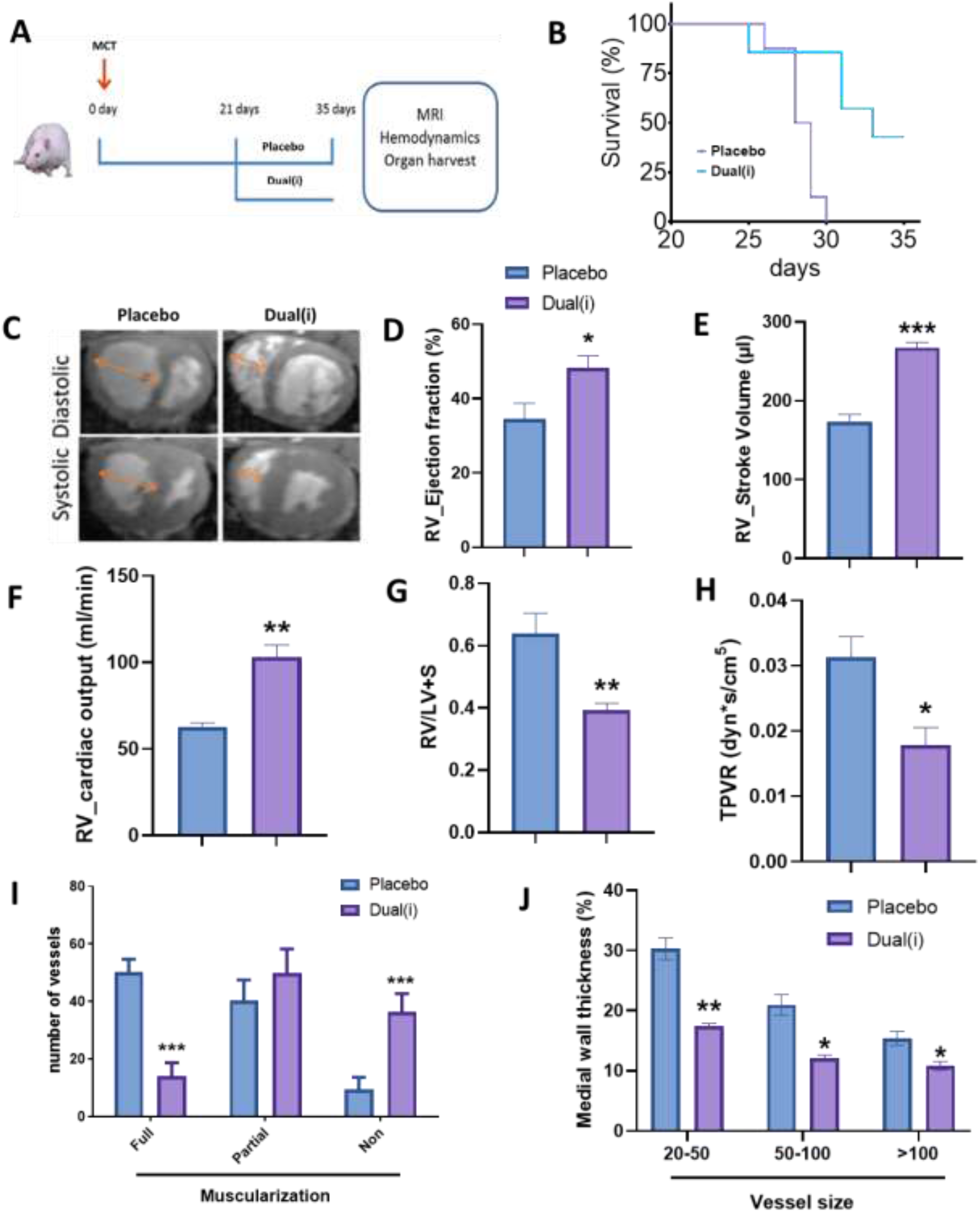
Effect of dual inhibitor on the MCT-PAH rat model. (A) Schematic representation of MCT rat model. (B) Survival curve and (C) Cardiac MRI images of MCT-PAH rats treated with placebo or dual inhibitor. (D) RV ejection fraction, (E) RV stroke volume, (F) cardiac output, and (G) Fulton index (RV+LV+S) from cardiac MRI from Placebo (n=6) and dual inhibitor (n=6) treated groups. Bars indicate means ± S.E.M. Data were analyzed using two-way ANOVA with Tukey’s multiple comparison. ***p <0.001 vs Placebo. (H) TPVR calculated from RVSP and Cardiac output of placebo (n=6) and Dual inhibitor treated groups (n=6). Bars indicate means ± S.E.M. Data were analyzed using the unpaired student t-test. *p <0.05 *vs* Placebo. (I) Medial wall thickness measurements in pulmonary vessels, (J) muscularization measurements from placebo (n=4) and Dual inhibitor treated groups (n=4). Bars indicate means ± S.E.M. Data were analyzed using two-way ANOVA with Tukey’s multiple comparison. *p <0.05, **p <0.01, ***p <0.001 *vs* Placebo.

**Figure 7.**
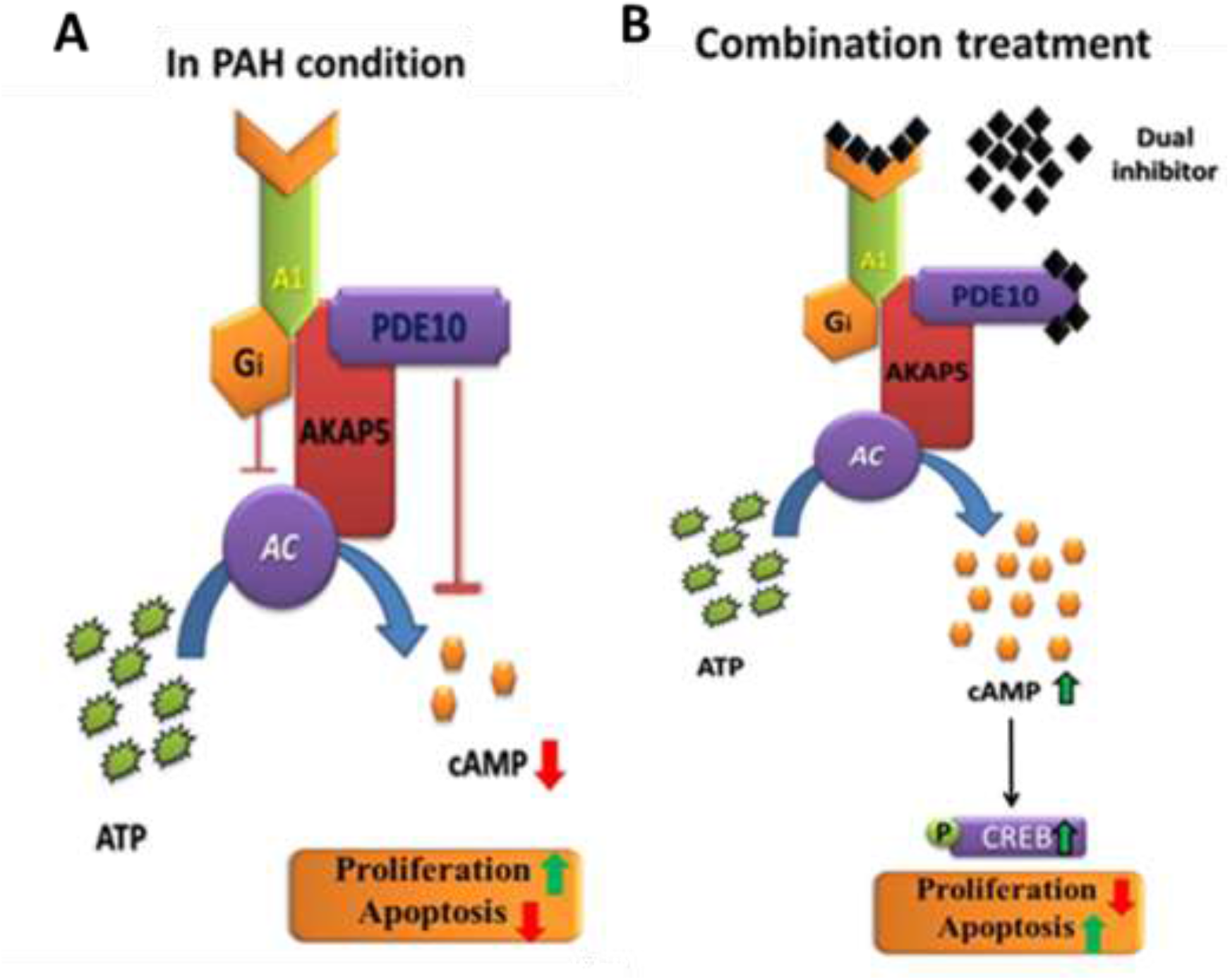
Schematic representation showing the regulation of cAMP by the ADORA1/PDE10A complex in IPAH condition (A) and effect mediated by the combination therapy in IPAH condition (B). A1-adenosine receptor A1, AC-adenylyl cyclase, ATP-adenosine triphosphate, cAMP-3’, 5’-cyclic adenosine monophosphate, PDE10A-phosphodiesteraste 10A, A-kinase anchoring protein 5, p-CREB-phosphorylated cAMP response element binding protein

## Discussion

In this study, we found for the first time that intracellular cAMP levels of pulmonary vascular smooth muscle cells are tightly regulated by ADORA1 and PDE10A, which form a regulatory complex exclusively under disease conditions. These distinct bidirectional ADORA1 and PDE10 regulatory complexes containing cAMP microdomains regulate cAMP dynamics in IPAH and promote the IPAH PASMC phenotype. In line, targeting both ADORA1 and PDE10A reversed the pathological phenotype and restored the decreased intracellular cAMP level in PASMCs of IPAH patients. Furthermore, the data from preclinical rat models of PAH highlight the therapeutic potential of the dual inhibitor of ADORA1 and PDE10A in PAH.

PAH is a multifactorial disease, characterized by increased pulmonary vascular pressure due to pathological remodeling and narrowing of the pulmonary vasculature, eventually leading to right heart failure and death. Previous studies have shown that secondary messengers such as cAMP play an important role in the maintenance of vascular smooth muscle cell quiescence and in the resolution of vascular remodelling via distinct repair mechanisms [27]. Previous studies from our lab has shown that IPAH patient-derived PASMCs exhibit pro-proliferative and anti-apoptotic feature [28-30], further in this study we verified the reduced intracellular cAMP levels in IPAH PASMCs. This finding is consistent with studies demonstrating impaired cAMP signalling in phenotypically altered vascular cells from PAH patients [23, 31-33].

The induction and inhibition of cAMP production is mediated by the activation of different GPCRs via different primary messengers, and the termination of cAMP signaling is mediated by phosphodiesterase enzymes (PDEs) via the hydrolysis of cAMP to 5′AMP [34, 35]. In PAH, PDE activity is increased under disease conditions. Apart from activity, the expression of several PDEs such as PDE1C, PDE3 and PDE10A is upregulated, and their inhibition has shown beneficial effects in PAH [23, 32]. Previous studies have also shown that targeting the prostacyclin receptor with prostacyclin analogs induces inhibition of proliferation in PASMCs and shows beneficial effects in PAH patients [36]. Similarly, adenosine, a potent vasodilator that also modulates intracellular cAMP production through receptor activation, has been shown to be reduced in the plasma of IPAH patients [15, 37].

The adenosine receptors (ADORA) are mainly of four types: the Gi-coupled ADORA1 and ADORA3, which inhibit cAMP production, and the Gs-coupled ADORA2a and ADORA2b, which induce cAMP production. In this study, we investigated the dysregulation of adenosine receptors under disease conditions and their potential as a therapeutic targets in PAH. The expression of ADORA1 was found to be strongly upregulated in IPAH patient-derived PASMCs and lung tissue compared to healthy samples. The upregulation of ADORA1 in both PASMCs and lung tissue from patients with idiopathic PAH (IPAH) is consistent with previous studies indicating altered expression of the adenosine receptor in cardiovascular disease [38-40]. Previously, targeting ADORA2b was shown to have beneficial effects in pulmonary hypertension associated with interstitial lung disease [20, 21]. The downregulation of intracellular cAMP levels in IPAH PASMCs observed in our study is consistent with the known inhibitory effect of ADORA1 on AC, resulting in reduced cAMP synthesis [41]. Silencing and inhibition of ADORA1 induced apoptosis and increased cAMP levels in healthy PASMCs, consistent with the known pro-apoptotic effects of increased cAMP levels [42]. However, the lack of a similar response in IPAH PASMCs suggests a disease-specific adaptation, possibly involving the formation of a more stringent cAMP-regulatory complex.

In agreement with our findings, a previous study showed that the induction of cAMP levels in IPAH patient-derived PASMCs was very low compared to healthy PASMCs when treated with forskolin (direct adenylyl cyclase activator) or beraprost (a PGI2 receptor agonist). Interestingly, the administration of forskolin or beraprost in combination with a general PDE inhibitor (IBMX) significantly increases intracellular cAMP levels in IPAH PASMCs was administered [32]. It was also previously reported that the expression of several cAMP-controlling PDEs is upregulated in PAH [43]. This motivated us to hypothesize that ADORA1 may be in close proximity to a cAMP-controlling PDE to form a bidirectional cAMP-regulatory complex under disease conditions. In our study, ADORA1 was indeed found to be in close proximity to PDE10A, a well-known cAMP-regulating PDE, exclusively in IPAH PASMCs. A previous study from our laboratory has shown that the expression of PDE10A is increased in IPAH compared to healthy individuals [23]. Moreover, other studies have shown that certain GPCRs and PDEs form complexes to stringently and spatially regulate cAMP in different biological contexts [44]. The study by Johnstone et. al. shows that PDE8 selectively regulates cAMP signaling through β2-adrenergic receptors and adenylyl cyclase 6 (AC6) in human airway smooth muscle cells, but not cAMP signaling mediated by the E-prostanoid receptor 2/4–AC2 complex [45]. In hepatocytes, the adenosine receptor A2a was found to form a complex with PDE3A and AC6 with the anchoring protein AKAP79 [46]. Similarly, our results show that in IPAH cells the supercomplex formed by ADORA1, PDE10A and AKAP5 crucially regulates cAMP at subcellular levels. Thus, we hypothesized that combined inhibition of ADORA1 and PDE10A could reverse the disease phenotype observed in IPAH PASMCs. Interestingly, siRNA-mediated silencing of both ADORA1 and PDE10A reduced proliferation and induced apoptosis in IPAH PASMCs, which was not previously observed with ADORA1 inhibition alone. The newly developed dual inhibitor targeting both ADORA1 and PDE10A also showed similar results to siRNA treatment in IPAH-PASMCs. Dual inhibition of ADORA1 and PDE10A resulted in a strong increase in intracellular cAMP levels and downstream phosphorylation of CREB. Similar to the results observed in this study, simultaneous activation of adenosine receptor 2A using an agonist and concomitant inhibition of PDEs such as PDE 2, 3, 4 and 7 induced cell death and intracellular cAMP levels in multiple myeloma (MM) and B-cell lymphoma cell lines [47]. Another study has shown that agonist-mediated activation of adenosine receptor A2a and concomitant inhibition of PDE10A has a potent anti-proliferative effect in non-small lung cancer cell lines [48]. Based on these solid in vitro data, the therapeutic potential of targeting ADORA1 and PDE10A with a dual inhibitor was investigated in monocrotaline (MCT) and Sugen5416-hypoxia (SuHx) rat PAH models. Strikingly, we found that treatment with the dual inhibitor targeting ADORA1-PDE10A improved survival in the rat MCT-PAH model. In both the MCT and SuHx-PAH models, the combination of ADORA1 and PDE10 inhibition improved cardiac parameters such as cardiac output, RV ejection fraction and stroke volume. Total pulmonary vascular resistance and pulmonary vascular remodeling were also reduced by the dual inhibition of ADORA1-PDE10 in both experimental PAH models. These results demonstrate that the combined inhibition of ADORA1/PDE10A by a dual inhibitor has therapeutic effects in vivo in the experimental PAH models.

The overall results of this study relate to the formation of a unique bidirectional cAMP-regulatory complex by ADORA1 and PDE10A under disease conditions. Therefore, we propose that targeting the ADORA1-PDE10A complex may provide a new therapeutic option for PH.

## Acknowledgments

We would like to acknowledge excellent technical contributions of Katharina Leib, Uta Eule, Natascha Wilker. Funding: This study was funded by the German Research Foundation (DFG), Collaborative Research Center (CRC) 1213 (Project A01, A05 grants to S.S.P., and A10N grant to R.S.) and European Research Council (ERC) Consolidator Grant (866051 to S.S.P.). R.S., W.S. and S.S.P. are supported by the Excellence Cluster Cardio Pulmonary Institute (CPI; EXC2026) and the German Center for Lung Research (DZL), and W.S., R.S. and S.S.P. are supported by the Max Planck Society.

