## Supplementary_Methods and tables for "Targeting ADORA-PDE10 cAMP Microdomain: A Novel Therapeutic Approach for Pulmonary Hypertension"

### Supplementary Material

### Methods

#### Human PASMCs isolation and culturing

The explanted lung tissues following lung transplantation from patients diagnosed with IPAH were obtained for the isolation of IPAH patient derived pulmonary artery smooth muscle cells (hPASMCs). The donor or healthy PASMCs were either obtained from Lonza (cat# CC-2581) or lung trimmings obtained from the downsized of healthy lung during transplantation. The study protocol for tissue donation was approved by the ethics committee (Ethik Kommission am Fachbereich Humanmedizin der Justus Liebig Universität Giessen) of the University Hospital Giessen (Giessen, Germany) in accordance with national law and with Good Clinical Practice/International Conference on Harmonisation guidelines. Written informed consent was obtained from each individual patient or the patient's next of kin (AZ 58/15). The pulmonary artery was isolated from the lungs and cut open to expose the luminal surface. The intima was scrapped out gently using a scalpel and the medial region was peeled away from the adventitial layer. The medial layer is cut into smaller sections and transferred to T75 flasks containing smooth muscle cells growth media (SmGM2, Lonza). Once the cells have migrated from the tissue and adhered to the plate, at 90% confluence the cells are passaged and further stored in the liquid nitrogen in the freezing media. The isolated PASMCs were always used for the experiments in the fifth or sixth passages.

#### RNA Isolation and qPCR

RNA was isolated from the cells and the tissues using the TRIzol™ reagent (Invitrogen) and concentration was measured using a nanodrop system. Equal amount of RNA was taken for the cDNA preparation using the cDNA synthesis kit (Thermoscientific) according to the manufacturer's instructions. mRNA expression analysis was performed using the qPCR experiment with the iQ SYBR green supermix kit (BioRad). The Primers used for the qPCR analysis was designed using the sequence information obtained from the Ensembl database and was purchased from Metabion (Martinsried, Germany). The ΔCt was calculated by normalizing the target gene to the house keeping genes ($\Delta Ct = Ct_{housekeeping} - Ct_{target}$). The graphs are represented as relative mRNA expression by normalizing the control sample set as 1. The Primers used in this study is listed in the [supplementary table 4].

#### Proteing Isolation and Western blotting

Protein is isolated using RIPA buffer (Thermoscientific) supplemented with protease inhibitor (Roche), after adding the protein isolation buffer the cells are scrapped and collected in the 1.5ml Eppendorf tube. The lysates are centrifuged at 10,000rpm for 20mins and the supernatant is transferred into a new tube and protein concentration was measured using DC protein assay (BioRad) according to manufacturer's instruction. The protein concentration is normalized and prepared in loading buffer. The protein is resolved using 7%, 10% or 12% sodium dodecyl sulfate (SDS)-polyacrylamide gel electrophoresis (depending on the size of the proteins). The proteins are blotted on to the PVDF membrane followed by probing using primary antibodies overnight at 4°C. The membranes are washed and probed with HRP conjugated secondary antibody, followed by development using the SuperSignal West Femto substrate (Thermoscientific) and imaged using ImageQuant LAS 4000 system (GE Healthcare). The blots were quantified using ImageJ, relative expression was calculated after normalization to loading controls (β-actin).

**Immunofluroscence Staining**

Isolated lungs were fixed in 4% PFA and embedded in paraffin. Sections, 3 µm thick, were prepared using a microtome and then heated at 65°C, followed by deparaffinization and rehydration with xylene and an ethanol series. Antigen retrieval was performed by heating in sodium citrate buffer (pH 6.0) for 15 minutes, followed by a 30-minute rest at room temperature (RT). The sections were permeabilized with 0.1% Triton in 1X PBS, then blocked with 10% BSA in PBST for 30 minutes. Specific primary antibodies [supplementary table 1]were added to the sections and incubated overnight at 4°C, followed by probing with fluorescent-tagged secondary antibodies [supplementary table 2] for 2 hours at room temperature. Nuclei were stained with DAPI and mounted with fluorescent mounting media (Dako). Imaging was performed on a Zeiss LSM700 confocal microscope using Zen Black software.

**Immunocytochemistry**

hPASMCs were seeded in eight-well chamber slides (15,000 cells/well) and cultured for 24 hours. Cells were then fixed in acetone: methanol (1:1) for 20 minutes at 4°C, washed in 1X PBS, and blocked (5% BSA, 5% goat serum, 0.1% Triton X-100 in PBS) for 1 hour at RT. Primary antibodies [supplementary table 1]diluted in blocking buffer were applied and incubated overnight at 4°C. Following three washes with 1X PBS, cells were incubated with Alexa flour secondary antibodies [supplementary table 2] diluted in blocking buffer for 2 hours at RT. After an additional three 1X PBS washes, cells were stained with DAPI for 5 minutes, and slides were mounted with anti-fade medium. Imaging was performed using a Zeiss LSM700 confocal microscope with Zen Black software.

**Intracellular cAMP concentration measurement**

Intracellular cAMP levels in hPASMCs were measured using a competitive non-acetylated EIA assay (Cayman Chemical) according to the manufacturer's protocol. hPASMCs were seeded in 6-well plates and treated with siRNA or inhibitors. After washing with 1X PBS, the cells were lysed in 0.1 M HCl for 15 minutes, and lysates were centrifuged at 12,000 rpm for 30 minutes at 4°C, with supernatants stored at -80°C. Standards and samples were added to anti-rabbit antibody-coated plates along with the cAMP tracer and antibody. The plates were incubated overnight at 4°C on a shaker. The next day, the wells were washed and treated with Ellman's reagent for 90 minutes at room temperature. Absorbance was measured at 412 nm, and free cAMP concentrations were derived from a standard curve using Magellan software. Results were normalized to total protein content, determined by the DC protein assay, and expressed as pmol of cAMP per mg (pmol/mg) of total protein.

**Cell proliferation Assay**

Cell proliferation in donor and IPAH PASMCs was quantitatively measured using a colorimetric BrdU incorporation assay kit (Roche), as per manufacturer's guidelines. For the experiments, hPASMCs were plated in 96-well plates at a density of 6,000 cells per well. After 24 hours, the cells were either transfected with siRNAs or treated with inhibitors. Once the treatment period was complete, cells were exposed to BrdU labeling solution for 2 hours. The media was then discarded, and the cells were fixed at room temperature for 30 minutes using 200 µl of FixDenat solution per well. After fixation, the cells were incubated with anti-BrdU-POD for 3 hours at room temperature. The solution was then discarded, and the wells were washed three times with wash buffer. Next, 100 µl of substrate solution was added to each well until colour development occurred, and absorbance was subsequently measured at 370 nm using a reference wavelength of 492 nm on an ELISA plate reader.

**Cell Apoptosis assay**

Cell death measurements in donor and IPAH hPASMCs were conducted using an in-situ cell death assay kit based on a quantitative sandwich enzyme immunoassay as per manufacturer's guidelines (Roche). For this assay, hPASMCs were cultured in 48-well plates, with 15,000 cells seeded per well. After 24 hours, the cells were either transfected with siRNAs or treated with inhibitors. Once the treatment period was complete, cells were centrifuged at 200g for 10 minutes and media was then discarded. 100 µl of 1X lysis buffer was added to each well and incubated at RT for 30 minutes. The plates were subsequently centrifuged at 300g for 10 minutes, and 40 µl of lysate was transferred into a streptavidin-coated microplate. An immunoreagent mix was prepared by combining anti-histone (biotin-labeled) and anti-DNA (HRP-labeled) antibodies in a 1:1:10 ratio with 1X incubation buffer, and 80 µl of this mix was pipetted into the microplate and incubated for 2 hours at RT with 300 rpm shaking. After incubation, the plates were washed three times with incubation buffer and then treated with 100 µl of substrate solution until color development. Finally, the absorbance of the plates was measured at 405 nm, with a reference wavelength of 409 nm, using an ELISA plate reader.

**siRNA treatment**

Knockdowns of ADORA1 and PDE10A were achieved using Dharmacon's siRNA smart pool technology [supplementary table 3], with Lipofectamine 3000 employed for transfection. A scrambled siRNA served as the negative control. hPASMCs were seeded at 150,000 cells per well in 6-well plates and cultured for 24 hours. The following day, the media was replaced with transfection media (50% Opti-MEM, 50% basal SMC media, 0.1% FCS), and the cells were incubated for 30 minutes. A transfection master mix was prepared with 250 µl of Opti-MEM, 1.5 µg of siRNA, and 7.5 µl of Lipofectamine 3000, which was then incubated for 30 minutes at room temperature before being added dropwise to the wells. After 6 hours, the media was exchanged for PASMC growth media. RNA was isolated 24 hours post-transfection, and protein was extracted after 48 hours. For proliferation and apoptosis assays, the cells were trypsinized, counted, and reseeded into 96-well and 48-well plates 24 hours post-transfection, with measurements taken 72 hours later.

**Inhibitor Treatment**

Inhibitor studies were conducted using an ADORA1 antagonist (10µM) and a novel dual inhibitor (3µM) developed by Palobiofarma (Barcelona) targeting both ADORA1 and PDE10A. Cells were seeded and cultured in PASMC growth media for 24 hours, followed by washing with DPBS and serum starvation in PASMC basal media with 0.1% FCS for another 24 hours. After serum starvation, the cells were treated with the inhibitors in PASMC growth media. Proliferation and apoptosis assays were performed 24 hours after treatment, cAMP studies were conducted 30 minutes post-treatment, and protein isolation was carried out after 6 and 24 hours of treatment.

**Co-immunoprecipitation**

Co-immunoprecipitation (Co-IP) was utilized to analyze protein-protein interactions. Cells were grown in 10 cm dishes, and lysis buffer was prepared by combining a protease inhibitor

(Roche) with Pierce IP-lysis buffer and 1 mM sodium orthovanadate. After incubating with lysis buffer, cells were scraped into 1.5 ml tubes and centrifuged at 13,000 rpm for 15 minutes at 4°C. The supernatant was collected, and protein concentration was measured using the Bio-Rad DC protein assay kit. For each condition, 500 µg of protein was diluted to 1 ml with IP-lysis buffer and pre-cleared with protein G-sepharose beads. The pre-cleared supernatant was incubated overnight at 4°C with antibodies against ADORA1 or AKAP5 or IgG controls. After overnight incubation, the G-sepharose beads were added for 3 hours to capture the immune complexes. The beads were washed five times with IP wash buffer (50 mM Tris-HCl (pH 7.4), 15 mM EDTA, 0.1% Triton X-100, and 100 mM NaCl). Following the washes, 50 µl of 2X sample buffer was added to the beads, which were then incubated at 95°C for 10 minutes. Western blotting was subsequently performed to analyze the immunoprecipitated proteins.

**Proximity ligation Assay**

The proximity ligation assay (PLA) was performed using the Duolink In Situ PLA kit (Sigma), based on proximity ligation and rolling circle amplification. hPASMCs were seeded in 8-well chamber slides and cultured for 24 hours. Cells were washed with 1X DPBS, fixed with ice-cold acetone: methanol (1:1) for 10 minutes at 4°C, and then blocked with 5% BSA in 1X PBST for 30 minutes. Primary antibodies were applied overnight at 4°C, followed by three washes with 1X DPBS. PLA probes (1:5) were incubated for 1 hour at 37°C, then treated with ligation reagent (1:5) for 30 minutes, and amplified with polymerase solution. Nuclear staining was performed with DAPI for 10 minutes, followed by washing and mounting with anti-fade medium. Images were acquired using a Zeiss LSM700 confocal microscope, detecting in the Texas-red range.

**In vivo PAH rat models**

Rats for this study were sourced from Charles River Laboratories (Sulzfeld, Germany), with all procedures following NIH guidelines. All the experiments were performed with permission from University Animal Care Committee and the Federal Authorities at Regierungspräsidium Darmstadt, Germany (approval no: V54-19c20/15-B2/1079). In vivo studies employed monocrotaline-induced and sugen hypoxia-induced pulmonary arterial hypertension models.

Monocrotaline Rat model

Sprague-Dawley (SD) rats were used to establish monocrotaline (MCT) model. To establish disease, 60 mg/kg of MCT was injected for 3 weeks, after which rats were randomized to receive either a placebo or treatment. The placebo group was given a nitrocellulose solution, while the treatment group received 10 mg/kg of the dual inhibitor (C21) by oral gavage. After

2 weeks of treatment, cardiac MRI and hemodynamic measurements were conducted for functional analysis, followed by organ harvesting for lung morphometric analysis.

Sugen hyoxia rat PH models

Wistar Kyoto rats were used to establish the Sugen hypoxia (SuHx) model. To induce disease, Su5416 (25 mg/mL in DMSO) was administered via subcutaneous injection, and rats were placed in a hypoxia chamber (10% $O_2$) for 3 weeks, followed by a return to normoxic conditions. Rats were then randomized to receive either a dual inhibitor (C21, 10 mg/kg) or a placebo via oral gavage for 2 weeks. For functional assessment, cardiac MRI and hemodynamic measurements were performed, followed by organ harvesting for lung morphometric analysis.

**Rat cardiac MRI**

Cardiac MRI measurements were performed on a 7.0 T Bruker Pharmascan (Bruker, Ettlingen, Germany), equipped with a 760 mT/m gradient system, using a room temperature 20 mm planar surface $^1$H receiver-coil and a 72 mm room temperature volume resonator for transmission and the IntraGate$^{TM}$ self-gating tool. The measurement were based on the gradient echo method (repetition time = 6.2 ms; echo time = 1.3 ms; flip angle = 10°; field of view = 45x45 mm; slice thickness = 1.0 mm; matrix = 128 x 128; oversampling = 100; number of frames = 20). The imaging plane was localized using scout images showing the 2- and 4-chamber view of the heart, followed by acquisition in short-axis view, orthogonal on the septum in both scouts. Multiple contiguous short-axis slices consisting of X to Y slices were acquired for complete coverage of the left and right ventricle. Rats were measured under volatile isoflurane (2.0 % in the air with a flow rate of 0.5 L/min) anesthesia; the body temperature was maintained at 37°C by a thermostatically regulated water flow system during the entire imaging protocol. MRI images are analyzed using MASS4Mice digital imaging software (Medis, Leiden, Netherlands).

**Hemodynamic measurements of rat**

Hemodynamic measurements were as previously described [28]. After the treatment duration, the rats were anesthetized by intraperitoneal injection of ketamine (9mg/kg body weight) and medetomidine (100µg/kg body weight), followed by intramuscular injection of heparin. A tracheotomy was performed on the anesthetized rats and was ventilated at 60 breaths/min frequency. A catheter (PE 50 tube) was inserted through the right jugular vein to measure the right ventricular pressure and the arterial pressure was measured by inserting the catheter into the left carotid artery. Following the hemodynamic measurement, the heart was harvested and the right ventricle (RV) and the left ventricle along with septum (LV+S) were separated and weighed, this was was used to calculate the Fulton index [RV/ (LV+S)].

**Lung vascular morphometric analysis**

Medial wall thickness of lung vasculature

To assess medial wall thickness (MWT), Weigert-Van Gieson staining was performed on 4 µm lung sections as previously described [28]. Sections were deparaffinized, rehydrated, and stained with Resorcin-Fuchsin solution overnight. After washing, slides were treated with Weigert's hematoxylin and Van Gieson solution, then dehydrated, mounted, and imaged using a Leica DM6000B microscope. Analysis was performed with Leica Qwin V3 software, and MWT was calculated as: %MWT = (2 × medial wall thickness / externalsss diameter) × 100.

Muscularization of Lung vasculature

To assess vascular muscularization, double immunohistochemistry was performed on 4 µm rat lung sections for α-SMA and von Willebrand factor, following methods described previously [28]. Sections were deparaffinized, rehydrated, and antigen retrieval was done using trypsin. Primary antibody staining for α-SMA and von Willebrand factor was conducted with blocking and color development using Vector and DAB substrates, respectively. Methyl green counterstaining was applied, followed by dehydration and mounting. Muscularization was quantified with the Leica DM6000B microscope and Leica Qwin V3 software.

**Statistical Analysis**

Results are reported as mean ± standard error of the mean (SEM). For statistical analysis, an unpaired Student's t-test was used for two-group comparisons, while one-way ANOVA with Dunnett's post hoc test was applied for multiple group comparisons. Statistical significance was set at $p < 0.05$. Exact group sizes (n) are indicated for each condition, representing independent biological replicates.

### List of tables

#### 1. List of Primary Antibodies

| Antibody Name | Technique | Source | Company |
| --- | --- | --- | --- |
| ADORA1 | WB/Co-IP | rabbit | Abcam |
| ADORA1 | IF | mouse | Santa Cruz Biotechnology |
| PDE10A | WB/Co-IP | goat | Santa Cruz Biotechnology |
| PDE10A | IF | rabbit | Abcam |
| AKAP5 | WB/Co-IP | mouse | BD Transduction Laboratories |
| PDE1 | IF | rabbit | Abcam |
| PDE3 | IF | rabbit | Abcam |
| PDE4 | IF | rabbit | Abcam |
| CREB | WB | rabbit | Cell Signaling Technology |
| P-CREB (Ser 133) | WB | rabbit | Cell Signaling Technology |
| a-SMA | IF | mouse | Sigma Aldrich |
| Von Willebrand factor | IF | mouse | Agilent |

WB: western blotting; IF: Immunofluorescence: Co-IP: Co-Immunoprecipitation

#### 2. List of Secondary Antibodies

| Antibody Name | Company |
| --- | --- |
| Alex Flour 488 anti-rabbit IgG | Life Technologies |
| Alex Flour 568 anti-rabbit IgG | Life Technologies |
| Alex Flour 488 anti-mouse IgG | Life Technologies |
| Alex Flour 568 anti-mouse IgG | Life Technologies |
| HRP conjugated anti-rabbit IgG | Sigma Aldrich |
| HRP conjugated anti-mouse IgG | Sigma Aldrich |
| HRP conjugated anti-Goat IgG | R&D systems |

#### 3. List of siRNAs

| siRNA | Sequence (5‘-3‘) | Company |
| --- | --- | --- |
| SCR_SMARTpool | 1) UGGUUUACAUGUCGACUAA | Dharmacon |

| | | |
|---|---|---|
| | 2) UGGUUUACAUGUUGUGUGA<br>3) UGGUUUACAUGUUUUCUGA<br>4) UGGUUUACAUGUUUUCCUA | |
| ADORA1_SMARTpool | 1) GGUAGGUGCUGGCCUCAAA<br>2) GGAGUCUGCUUGUCUUAGA<br>3) CAAGAUCCCUCUCCGGUAC<br>4) AUCCAGAAGUUCCGCGUCA | Dharmacon |
| PDE10A_SMARTpool | 1) GAACUAAACAGCUAUAUAG<br>2) GUAAUUGGUUUGAUGAUGA<br>3) GAACAAGGAGUUAUAUUCA<br>4) GCAGAGGCCUUGCCAAACA | Dharmacon |

#### 4. List of primers

| **Name** | **Species** | **Sequence (5' – 3')** |
|---|---|---|
| ADORA1 | Human | F: GGCAGGTGAGGAAGGGTTTA<br>R: CTGCCTGACTGTTCTGTCCA |
| ADORA2a | Human | F: GCGCAGTATGAGAGGGCTTA<br>R: GAGGCACCCCAGCACTATC |
| ADORA2b | Human | F: ATACCTGGCCATCTGTGTCC<br>R: GAGTCAATCCGATGCCAAAG |
| PBGD | Human | F: CAGGAGTCAGACTGTAGG<br>R: ACTCTCATCTTTGGGCTGTTTTC |
| PDE10A | Human | F: CTCCGACATGGAAGATGGAC<br>R: CCCCCATCTCTTCAACTGTAA |
| B2M | Human | F: AGATGAGTATGCCTGCCGTG<br>R: TCATCCAATCCAAATGCGGC |
| AKAP5 | Human | F: GACTGCAGCATCAAAGTCCA<br>R: CATCTTTCCGTGAGATGCCT |
